# Assessing chromatin relocalization in 3D using the patient rule induction method

**DOI:** 10.1101/2021.05.08.443243

**Authors:** Mark R. Segal

**Affiliations:** Department of Epidemiology and Biostatistics, University of California, San Francisco CA 94143-0560 USA

## Abstract

Three dimensional (3D) genome architecture is critical for numerous cellular processes, including transcription, while certain conformation-driven structural alterations are frequently oncogenic. Inferring 3D chromatin configurations has been advanced by the emergence of chromatin conformation capture assays, notably Hi-C, and attendant 3D *reconstruction* algorithms. These have enhanced understanding of chromatin spatial organization and afforded numerous downstream biological insights. Until recently, *comparisons* of 3D reconstructions between conditions and/or cell types were limited to prescribed structural features. However, *multiMDS*, a pioneering approach developed by Rieber and Mahony (2019) that performs joint reconstruction and alignment, enables quantification of all locus-specific differences between paired Hi-C data sets. By subsequently mapping these differences to the linear (1D) genome the identification of *relocalization* regions is facilitated through use of peak calling in conjunction with continuous wavelet transformation. Here, we seek to refine this approach by performing the search for significant relocalization regions in terms of the 3D structures themselves, thereby retaining the benefits of 3D reconstruction and avoiding limitations associated with the 1D perspective. The search for (extreme) relocalization regions is conducted using the *patient rule induction method* (PRIM). Considerations surrounding orienting structures with respect to compartmental and principal component axes are discussed, as are approaches to inference and reconstruction accuracy assessment. Illustration makes recourse to comparisons between four different cell types.

## 1 Introduction

The three-dimensional (3D) organization of chromatin within the eukaryotic nucleus is central to numerous biological processes, including transcription, replication, development and even memory formation (Marco *and others* (2020)). It follows that organizational *changes* are also consequential, with select perturbations being oncogenic (Flavahan *and others* (2016)), while architecture impacting mutations can disrupt gene expression resulting in developmental disorders and disease (Krijger and de Laat (2016); Spielmann *and others* (2018)). Accordingly, identifying such changes is fundamental to understanding of evolutionary and regulatory mechanisms.

Just as much of the current understanding of global principles of hierarchical chromatin organization derives from Hi-C and related assays (Lieberman-Aiden *and others* (2009); Dixon *and others* (2012); Rao *and others* (2014); Bonev and Cavalli (2016)), so too has study of chromatin changes focused on attendant data types, notably contact maps or matrices. A variety of approaches investigating associated dynamics have been proposed, including (i) visual comparisons (Crane *and others* (2015); Darrow *and others* (2016)), (ii) global measures of contact map agreement (Yang *and others* (2017); Yan *and others* (2017)), (iii) targeted comparisons of pre-defined features, in particular topologically associating domains (TADs) (Huynh and Hormozdiari (2019)) and loops (Rao *and others* (2014); Lareau and Aryee (2018)), and (iv) identifying pairs of genomic loci (contact matrix bins) with significantly differential counts (Lun and Smyth (2015); Djekidel *and others* (2018)). Each of these classes has readily identified shortcomings, as detailed in recent work that advances a compelling algorithm (CHESS; Galan *and others* (2020)) for the comparison of chromatin contact maps and automatic differential feature extraction.

All these methods operate strictly on the contact map level. However, there have several illustrations of benefits in proceeding from a contact map to an inferred 3D reconstruction, this added value deriving from the ability to superpose genomic attributes on the reconstruction. Examples include co-localization of genomic landmarks such as early replication origins in yeast (Witten and Noble (2012); Capurso and Segal (2014)), gene expression gradients in relation to telomeric distance and co-localization of virulence genes in the malaria parasite (Ay *and others* (2014)), the impact of spatial organization on double strand break repair (Lee *and others* (2016)), and elucidation of ‘3D hotspots’ corresponding to (say) overlaid ChIP-Seq transcription factor extremes which can reveal novel regulatory interactions (Capurso *and others* (2016)).

Correspondingly, we might anticipate that studying chromatin change would also benefit from assessments utilizing 3D reconstructions. That this is indeed the case was convincingly demonstrated in pioneering work of Rieber and Mahony (2019), hereafter RM. After devising an algorithm, multiMDS, reviewed in Section 2.2, that simultaneously obtains 3D reconstructions and alignments for paired Hi-C datasets, they proceed to demonstrate how the Euclidean distance between corresponding loci on the reconstructions, termed *relocalization* distance can be further analyzed, leading to a variety of biological insights. For example, by mapping relocalizations to genomic coordinates – the 1D (linear) genome – they were able to recapitulate known galactose-dependent re-organization of yeast genes, as well as identifying novel changes. Importantly, these findings would not have been elucidated using either independent 3D reconstruction or Hi-C loop calling. RM further investigated cell type associated changes that we revisit in Section 3.4.

The multiMDS algorithm itself operates by “borrowing strength” between related configurations with a tuning parameter governing the extent of borrowing. A related approach is taken by SIMBA3D ((Rosenthal *and others*, 2019)) which weighs reconstructions from single-cell Hi-C data toward a bulk cell solution. While we describe potential enhancements to multiMDS our main objective is to provide an alternative analytic approach to the resultant relocalization distances. Instead of mapping these back to the linear genome and discerning 1D *segments* of interest via peak finding methods we seek 3D *regions* of significant displacement. This is accomplished by superposing relocalization values on the multiMDS-inferred 3D reconstruction, then using the patient rule induction method (PRIM, Friedman and Fisher (1999); Hastie *and others* (2009)) to elicit focal extremes.

Our methodological development and subsequent results, including discussion of coordinate systems and axes, are presented in Sections 2 and 3 respectively. As illustrated, we believe that assessing relocalization in 3D confers advantages, concomitant with the motivation underscoring 3D reconstruction itself: the spatial organization of the folded genome is consequential and 1D representations can be limiting. Nonetheless, as detailed in the Discussion (Section 4) there are instances where our 3D approach is inapplicable and/or where improvements could be effected.

## 2 Methods

We position the development of our 3D approach to relocalization by initially briefly describing the problem of chromatin 3D reconstruction based on Hi-C data (Section 2.1), followed by an overview of the multiMDS method for *joint* reconstruction and alignment of paired data (Section 2.2). These aligned structures provide the basis for obtaining 1D relocalization scores. Notable relocalization segments, that are contiguous in genomic coordinates, are then identified using peak finding after continuous wavelet transformation, which is also briefly described (Section 2.3). Reconstructions derived from Hi-C data are inherently coordinate system free: in section 2.5 we detail two relevant orientation schemes, compartmental and principal axes. The latter has bearing on our 3D approach to relocalization that deploys PRIM which, along with permutation-based methods for inferring significant relocalized regions, is detailed in Section 2.4.

### 2.1 3D chromatin reconstructions from Hi-C data

We restrict attention to reconstruction of *individual* chromosomes; whole genome architecture can follow by appropriately positioning these solutions (Segal and Bengtsson (2015); Rieber and Mahony (2017)). As is standard, we disregard complexities deriving from chromosome pairing arising in diploid cells (which can be disentangled at high resolutions (Rao *and others* (2014)) or by imposing additional constraints (Belyaeva *and others* (2021)), as well as concerns surrounding bulk cell experiments and inter-cell variation.

The result of a Hi-C experiment is the *contact map*, a symmetric matrix 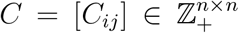 of contact counts between *n* (binned) genomic loci *i, j* on a genome-wide basis. Various approaches to contact matrix normalization have been proposed; our methods are agnostic to these. The 3D chromatin reconstruction problem is to use the contact matrix *C* to obtain a 3D point configuration *x*_1_, …, *x*_*n*_ *∈*ℝ^3^ corresponding to the spatial coordinates of loci 1, …, *n* respectively.

Many algorithms have been proposed to tackle this problem with broad distinction between optimization and model-based methods (Varoquaux *and others* (2014); Rieber and Mahony (2017)). A common first step is conversion of the contact matrix into a *distance* matrix *D* = [*D*_*ij*_] (Duan *and others* (2010); Varoquaux *and others* (2014); Ay *and others* (2014); Shavit *and others* (2014); Rieber and Mahony (2017)), followed by solving the *multi-dimensional scaling* (MDS; Hastie *and others* (2009)) problem: position points (corresponding to genomic loci) in 3D so that the resultant interpoint distances “best conform” to the distance matrix.

A variety of methods have also been used for transforming contacts to distances. At one extreme, in terms of imposing biological assumptions, are methods that relate observed intra-chromosomal contacts to genomic distances and then ascribe *physical* distances based on organism specific findings on chromatin packing (Duan *and others* (2010)) or relationships between genomic and physical distances for crumpled polymers (Ay *and others* (2014)). Such distances inform the subsequent optimization step as they permit incorporation of known biological constraints that can be expressed in terms of physical separation. However, obtaining physical distances requires both strong assumptions and organism specific data (Fudenberg and Mirny (2012)). More broadly, a number of approaches (Zhang *and others* (2013); Varoquaux *and others* (2014); Zou *and others* (2016); Rieber and Mahony (2017)) utilize power law transfer functions to map contacts to (non-physical) distances *D*_*ij*_ = (*C*_*ij*_)^*−α*^ if *C*_*ij*_ *>* 0

MDS operationalizes the above notion of “best conforms” via an objective function termed the *stress*, a standard version of which is:

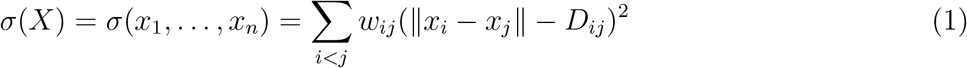

where *‖·‖* denotes the Euclidean norm and *w*_*ij*_ are analogous to precision weights often taken as 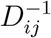 (Zhang *and others* (2013)) or 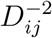 (Varoquaux *and others* (2014)). More elaborate variants incorporate penalties to ensure loci with *C*_*ij*_ = 0 are not positioned too close (Zhang *and others* (2013)). Several 3D reconstruction approaches use MDS as a building block (Duan *and others* (2010); Ay *and others* (2014); Shavit *and others* (2014); Segal and Bengtsson (2015); Rieber and Mahony (2017)), overlaying, for example, probabilistic modeling of contact counts (Varoquaux *and others* (2014); Tuzhilina *and others* (2020)), with the latter further imposing that the solution be a 1D contiguous curve in 3D via principal curve techniques (Hastie and Stuetzle (1989)).

### 2.2 MultiMDS

Our interest is in detecting pairwise structural differences between chromatin configurations corresponding to differing conditions or, as analyzed subsequently, differing cell-types. RM’s multiMDS algorithm is expressly designed for such detection, enabled by simultaneously inferring and aligning the two 3D structures. As they demonstrate, such a joint approach confers clear advantages over independently reconstructing the two configurations. Since multiMDS serves to underpin our relocalization approach we provide an overview, framed in terms of comparing cell types, noting aspects where further refinement is possible.

For a given chromosome from each cell type, we commence with two *n ×n* Hi-C contact matrices, *C, C′*, obtained using the same binning of genomic coordinates, then transformed to respective distance matrices *D, D′. We* seek the two corresponding 3D configurations 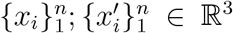. Central to multiMDS is minimization of an expanded MDS stress objective:

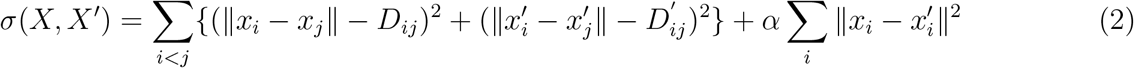

This criterion augments the (unweighted) individual stresses with a term that measures the sum of squared distances between corresponding points on the two solution configurations. The parameter *α* is termed the *similarity* weight and plays an important role: borrowing strength by minimizing the difference between the two (related) configurations. That this strategy is effective versus independent reconstruction (*α* = 0) was shown, in part, by comparing reconstructions using the same dataset. In this setting the root mean square distance (RMSD) between reconstructions should be zero, but was much greater for independent MDS than for multiMDS. Determination of the similarity weight is based on identifying the *α* value beyond which further increases do not improve reproducibility, as measured by the correlation between multiple runs with the same Hi-C data but differing random starting configurations. The algorithm proceeds by alternating updates of *X* and *X*^*′*^ that minimize the stress (2) using the previously devised *miniMDS* algorithm (Rieber and Mahony (2017)).

At the cost of increased computation several extensions to multiMDS are possible. First, the use of a global similarity weight is equivalent to using loci (bin) specific weights but prescribing that these are all equal. If there are a priori reasons for emphasizing particular loci then this could be *accommodated by allowing differential weighting. Second, following each cycle of X, X′* updates, Procrustes alignment (Hastie *and others* (2009)) of the respective structures could be applied. Third, rather than obtaining distance matrices using a prespecified value of the power law index, this index can be treated as a parameter and estimated via grid search (Zhang *and others* (2013)).

Finally, akin to (1), the individual stress components could incorporate precision weights. Here, our goal is comparison of current 1D approaches to relocalization with our proposed 3D approach and so, accordingly, we use the existing multiMDS scheme.

### 2.3 1D relocalization – continuous wavelet transform

On convergence, multiMDS provides 3D configuration estimates 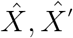 and the relocalzation (Euclidean) distance, at each locus *i*, between them: 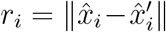. As described next, it is by analyzing *r* as a function of corresponding genomic coordinates that RM seek to probe relocalization features. However, as noted in the Introduction, this 1D viewpoint potentially discards the 3D context that motivated reconstruction to begin with.

Natural relocalization features to elicit are focal extremes, these representing the most pronounced structural differences. Using the 1D lens RM identify such relocalization *peaks* via the continuous wavelet transform (CWT; Du *and others* (2006)) applied to *r*. The CWT is given by

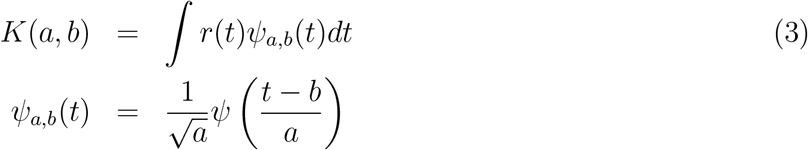

where *ψ*(*t*) is the *mother wavelet, a >* 0 is the scale, *b* is the translation, and *K* is the 2D matrix of wavelet coefficients. Throughout we take *ψ* as the Mexican Hat wavelet which is proportional to the second derivative of the Gaussian probability density function.

A heatmap depiction of *K* is provided in Figure 1. As detailed in Du *and others* (2006), finding peaks in *r* corresponds to finding ridges in this 2D CWT coefficient matrix. This approach is implemented in their companion R package MassSpecWavelet which further provides methods for filtering peaks based on signal-to-noise ratios (SNR). Here we adopt the default, yet stingent, peak calling criterion of SNR = 3.0. As the package name suggests, the CWT approach is geared toward mass spectrometry (MS) applications. But, the methodology for both peak identification and SNR attribution pertain generally, with the following caveats. MS spectra are obtained at equi-spaced (mass-charge) values; this aspect is used implicitly in CWT computation. For the most part, this is not problematic for Hi-C data since binning is into equi-spaced genomic intervals. However, centromeric regions, for example, are often absent from contact matrices due to mappability issues. In our applications both the extent of non equi-spaced genomic bins and the impact of linear interpolation to achieve equi-spacing is minimal. The Mexican Hat and, more generally, symmetric mother wavelets, was deemed appropriate for MS spectra without pre-processing based on baseline behavior. Whether these characteristics pertain to relocalization distances is uncertain and beyond the scope of the present work.

**Figure 1:**
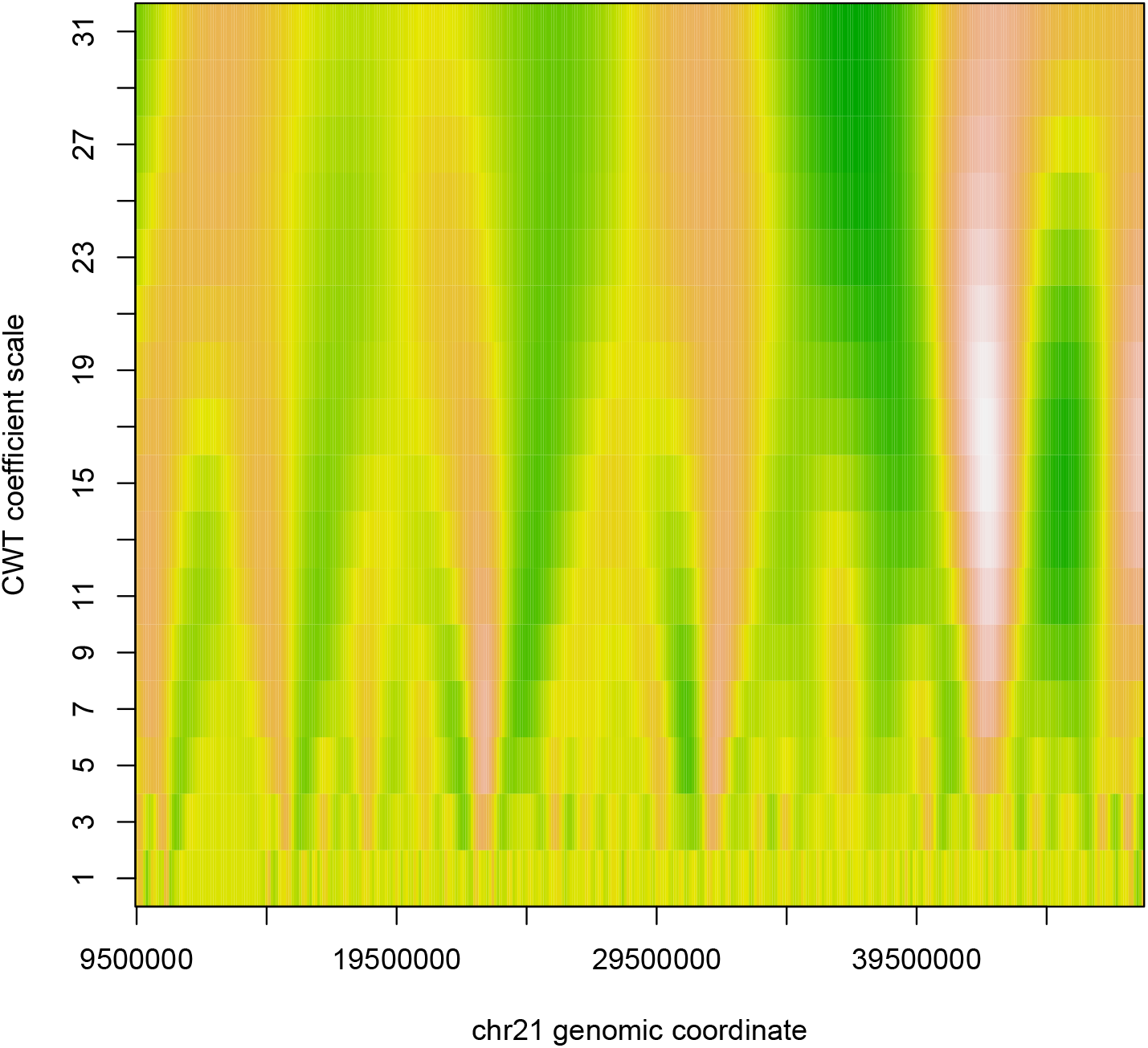
CWT coefficient image plot for IMR90 - HMEC relocalizations with yellow representing high amplitudes and green representing low amplitudes.

### 2.4 3D relocalization – patient rule induction method

CWT peak finding identifies focally extreme relocalizations with respect to genomic coordinates. Yet, the folding and compaction of chromatin means that loci that are close in 1D genomic coordinates may be far apart in 3D and vice versa, contiguity notwithstanding. Accordingly, it is purposeful to seek focally extreme relocalization regions in 3D with respect to the inferred structures themselves. We pursue this objective using the patient rule induction method (PRIM, Friedman and Fisher (1999)), as implemented in the R package `prim` (Duong (2020)).

The data set-up is as follows. As per Section 2.2, multiMDS provides aligned 3D reconstructions 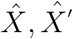 corresponding to the two conditions being contrasted. We arbitrarily select the first of these as referent with results being largely insensitive to this choice. For each genomic locus (bin), *i*, its three coordinate values 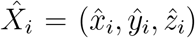 then serve as covariate values, while the relocalization distance *r*_*i*_ constitutes the outcome.

PRIM seeks to identify focally extreme outcomes (which here will be our relocalization distances) by sequentially and strategically paring away data regions (constrained to be slices) so that the *mean* outcome over the remaining data is increased. Instead of optimizing with respect to mean outcome alternate functions can be deployed, as commented on in the Discussion. At each iteration, a fraction `peel_alpha` of the loci are *peeled* off the 3D reconstruction by evaluating the extremal slices orthogonal to each of the reconstruction’s coordinate axes and removing loci from whichever slice results in the highest mean relocalization distance for the remaining loci. So, six candidate slices, corresponding to high and low values for each of the three coordinate axis, are evaluated per iteration. Hastie *and others* (2009, Figure 9.7) provide an illustrative cartoon for 2D (https://web.stanford.edu/~hastie/ElemStatLearn/printings/ESLII_print12_toc.pdf). This process is continued until a prescribed minimum number of loci `mass_min` remains. The resultant region can be enlarged to correct potential overshoot by a reverse *pasting* process utilizing a tuning parameter `paste_alpha`, set smaller than `peel_alpha`, if that increases the mean relocalization distance. At this point, a PRIM region or box has been identified, which represents a focal relocalization. The loci comprising this box are then excluded and the entire procedure is recursed to identify additional boxes.

As with clustering techniques, PRIM will produce boxes irrespective of relocalization signal. Accordingly, inferential techniques are necessary. Here we make recourse to permutation by repeatedly scrambling relocation distances over reconstruction loci, applying PRIM to each such permutation, and determining mean relocation distance for the resultant boxes. This provides a null referent distribution for assessing the actual box means obtained from the original data. It is worth noting that initial PRIM box elicitation, and subsequent permutation-based assessment thereof, are invariant to monotone transformations of relocation distances.

When it comes to characterizing significant boxes, akin to CWT analyses, we still reference the genomic coordinates involved. However, as emphasized, these relocalized regions may not necessarily be limited to consecutive coordinates but can reflect chromatin folding.

### 2.5 Compartment and principal component scores and axes

Hi-C based 3D chromatin reconstructions are intrinsically coordinate system free: there is no preferred orientation and rotated structures are equally valid. But, the PRIM procedure for identifying regions (boxes) with extreme relocalizations is conditional on the reconstruction’s (arbitrary) co-ordinate system since it is with respect to these axes that slices are defined. Here, as previously ((Capurso *and others*, 2016)), we counter this arbitrariness via sensitivity analyses involing applying PRIM to several randomly rotated structures.

Alternatively, coordinate system ambiguity can be resolved by rotating the 3D reconstruction to principal component (PC) axes prior to application of PRIM. Importantly, this approach does more than just provide a well-defined reference frame: there have been empiric and theoretic demonstrations that such rotation improves PRIM performance ((Dalal *and others*, 2013; Diaz-Pachon *and others*, 2017)). So throughout we apply PC rotation to 3D chromatin structures before using PRIM. Further justification for this approach derives from the (empiric) relationship between the first PC axis and another important aspect of chromatin organization described next.

That chromatin is organized into *compartments*, designated A and B, was proposed in the original Hi-C paper ((Lieberman-Aiden *and others*, 2009)) by way of explaining the plaid patterning of contact matrices. This patterning entails that chromatin interactions occur predominantly within, as opposed to between, compartments. Moreover, the compartments differentiate between active (A) and inactive or repressive (B) chromatin.

Compartments are defined as follows. A normalized contact matrix 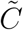 was obtained via 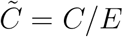, where operations are elementwise and *E* is the banded matrix of expected counts according to common genomic distance (a primary determinant of contact counts) and computed here as the mean of the diagonals of *C*. Principal components analysis (PCA) is applied to the correlation matrix of 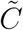 and scores of the first principal component (PC1) are termed *compartment scores*, with their sign determining compartment membership. Compartment scores strongly correlate with locus position along a physical axis between active nuclear interior and inactive lamina-associated domains (Luperchio *and others* (2017)). RM obtain this *compartment axis* in 3D by support vector regression (SVR) of compartment scores on their multiMDS 3D reconstruction coordinates. They contend that the strength of association between compartment scores and the reconstruction derived axis affirms that multiMDS is capturing consistent features of nuclear organization, rather than superficial similarities between the aligned structures. Further, they distinguish within and between compartmental relocalizations.

In Section 3.2 we establish (empiric) relationships between compartmental and principal component axes, the latter being important in the PRIM context as described above. While PCA is also utilized in generating compartment scores, and hence the attendant compartmental axis, there is no analytical connection between the two axes. So, in opting to use the 3D reconstruction PC axes as the coordinate system for our PRIM analyses we inherit – in view of the empiric correspondence – the interpretability of the compartment axis, but avoid issues and uncertainties (for example with defining expected counts and SVR tuning) in its estimation.

Each member of an aligned 3D reconstruction pair will have corresponding compartment scores obtained from its underlying contact matrix as described above. RM use absolute differences in these scores to further explore relocalizations that arise within versus between compartment types, although there is arbitrariness in thresholding these differences to obtain this dichotomy. Another concern is that compartment score differences are only meaningful in the face of sign ambiguity of the scores (due to sign ambiguity of the defining eigenvector) if this ambiguity is resolved, using external data. We investigated relationships between relocalization and compartments by (i) graphical overlay of significant relocalization regions and compartment score differences, and (ii) regressing relocalization distances on compartment score differences, then using residuals from these regressions as the response in PRIM- and CWT-region or segment identification efforts. However, no clear associations emerged from these analyses.

### 2.6 Reconstruction accuracy assessment

In order for any identified relocalization regions or segments to be interpretable it is essential that the underlying reconstructions from which relocalization distances are derived be accurate. However, even beyond definitional issues pertaining to bulk cell assays and diploid genomes, the absence of gold standards makes assessing accuracy challenging. While some comparisons of competing reconstruction approaches have made recourse to simulation (Zhang *and others* (2013); Varoquaux *and others* (2014); Zou *and others* (2016); Park and Lin (2017)), real data referents are preferable. Accordingly, appeal to fluorescence in situ hybridization (FISH) imaging is common. This proceeds by comparing distances between imaged probes with corresponding reconstruction-based distances. But such strategies are severely limited by the sparse number of probes (*∼*2 *−*6; see Lieberman-Aiden *and others* (2009); Shavit *and others* (2014); Park and Lin (2017)) and the modest resolution thereof, many straddling over 1 megabase (Mb). The recent advent of *multiplex* FISH (Wang *and others* (2016); Su *and others* (2020)) transforms 3D genome reconstruction accuracy evaluation by providing large numbers of replicate images possessing an order of magnitude more probes and hence two orders of magnitude more inter-probe distances than conventional FISH. Moreover, the probes are at higher resolution and centered at topologically associated domains (TADs; see Dixon *and others* (2012)). We use this imaging data, along with companion accuracy assessment approaches (Segal and Bengtsson (2018); Tuzhilina *and others* (2020)) to evaluate multiMDS reconstructions, although being restricted to the only cell line (IMR90) for which multiplex FISH data is available.

Briefly, we employ a three-step assessment procedure as follows. First, a *medoid replicate* is defined as the imaging replicate whose Procrustes dissimilarity to all other replicates is minimal – this 3D structure serves as our proxy gold standard. Second, we obtain an empirical reference distribution capturing experimental variation by measuring Procrustes dissimilarities between individual imaging replicates and the gold standard. Third, we align the multiMDS 3D reconstruction with the gold standard which involves coarsening of the reconstruction to yield comparable resolution. Quantification of how close this reconstruction is to the gold standard is then measured via dissimilarity and superposed on the referent distribution.

## 3 Results

### 3.1 Data sources and selections

Raw Hi-C counts were obtained for the IMR90 (embryonic lung fibroblast), HMEC (mammary epithelial), HUVEC (umbilical vein endothelium) and K562 (chronic myelogenous leukemia) cell lines from the Gene Expression Omnibus (accession GSE63525; (Rao *and others*, 2014)). Pairwise aligned 3D reconstructions between these cell lines, as well as for contrasting yeast conditions, obtained using multiMDS, were provided by Lila Rieber and Shaun Mahony. In the Discussion we detail why the yeast data were not amenable to PRIM-based analysis. The paired alignments consist of 3D coordinates at each genomic locus (bin) of the underlying contact matrix. The resolution (bin extent) of the aligned reconstructions was 100 kilobases (kb). Relocalization (distance) is then simply the Euclidean distance between corresponding points of the 3D structures.

Chromosome 21 was highlighted by RM since relocalizations among IMR90, HMEC and HUVEC pairs at 47.447.5 Mb were putatively identified and deemed noteworthy since the locus did not significantly differ in compartment score. This locus contains an intergenic segment featuring different histone modifications across these cell lines that potentially underscores the relocalizations. Moreover, a higher resolution analysis revealed differing degrees of close contact between relocalization segments within this locus. As a negative control RM also investigate an aligned multiMDS 3D reconstruction between K562-HUVEC since these cells have a similar epigenetic profile for the locus in question. We revisit locus identification and segment contact from the 3D relocalization region persepective in Section 3.4, as well as examining a variety of histone modifications using data from ENCODE, obtained as detailed in Table S2 of the supplementary material.

### 3.2 Compartmental and principal axes correspondence

In identifying focal relocalizations in the four human cell line pairs using CWT, RM emphasize genomic segments harboring relocalization peaks that overlap with segments featuring low (absolute) compartment score differences, which highlight *intra*-compartmental relocalizations. These are deemed of particular interest since inter-compartmental relocalizations were determined to be more commonplace based on the extent of components of the relocalization vector that project onto the compartment axis versus onto orthogonal axes, based on visual inspection. We note, however, that for these four pairs there is minimal overlap between CWT-identified relocalization peaks and differential compartment score peaks.

For our PRIM-based technique for detecting relocalization regions we take an indirect approach to using compartment scores and axes. This commences with performing PCA on the coordinates of each member of an aligned structure pair and obtaining PC1 scores. The (absolute) correlations between these PC1 scores and compartment scores obtained from the Hi-C contact matrix of the corresponding cell line are presented in Supplementary Table S1. Given the large numbers of constituent scores, which corresponds to the number of genomic bins underlying the reconstructions, the correlations are highly significant. Accordingly, in pursuing PRIM analyses on the PCA rotated coordinate system as recommended (Dalal *and others* (2013); Diaz-Pachon *and others* (2017)), we enjoy the added benefit of interpretability, since the first PC closely recapitulates the compartmental axis which, in turn, corresponds to the abovementioned physical axis between active nuclear interior and inactive lamina-associated domains. While this principal axis system underlies our primary analyses, in Section S3 of the supplementary material we showcase sensitivity analyses based on randomly rotated configurations, results being largely invariant.

### 3.3 IMR90 multiMDS reconstruction accuracy

Results from applying our accuracy assessment scheme (Section 2.6) to the multiMDS reconstruction of IMR90 chromosome 21 aligned to HMEC are shown in Figure 2 – almost identical findings pertain to IMR90 aligned to HUVEC. The histogram displays Procrustes dissimilarities between the mediod multiplex FISH image and each of the 111 multiplex FISH replicate images. That the multiMDS reconstruction is in the left tail of the distribution indicates that the reconstruction conforms with the inferred gold standard, akin to the PoisMS (Tuzhilina *and others* (2020)) but unlike the HSA (Zou *and others* (2016)) reconstructions.

**Figure 2:**
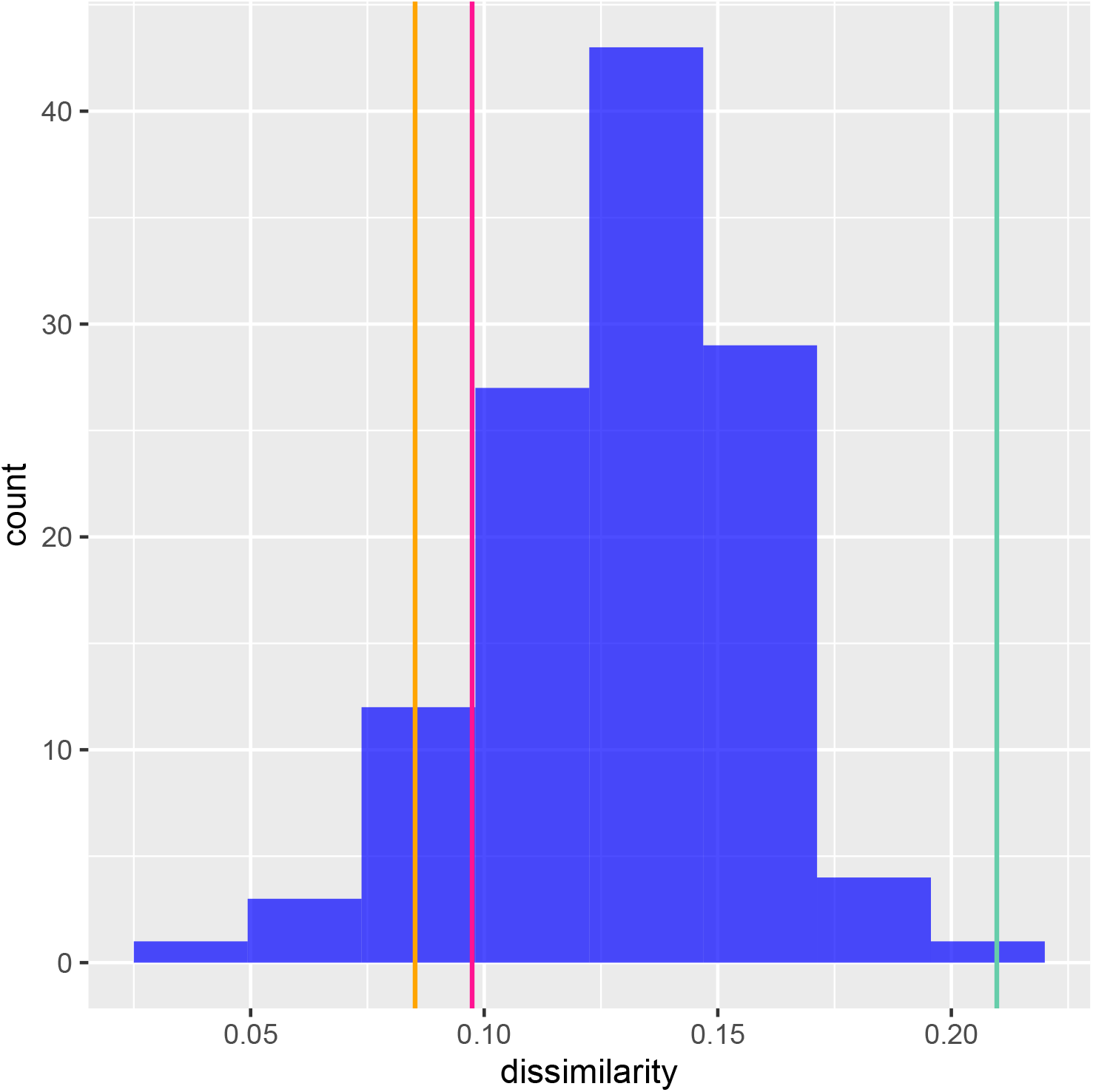
Reference distribution (blue histogram) depicting the dissimilarity between the inferred gold standard and 111 multiplex FISH replicate conformations IMR90 for chromosome 21. The vertical lines correspond to dissimilarities between the gold standard and select 3D reconstructions: red – multiMDS (aligned to HMEC), orange – PoisMS (Tuzhilina *and others* (2020)), green – HSA (Zou *and others* (2016)).

### 3.4 PRIM and CWT identified relocalization regions

We present results pertaining to identification of notable relocalization regions and segments for the four pairwise cell line comparisons via a series of figures. In Figure 3 we display histograms of the permutation distribution – obtained by permuting (1000 times) the original relocalization values over the aligned structure coordinates – of the top three PRIM-selected relocalization region (box) means for the IMR90 - HMEC cell line comparison. Supplementary Figures S1-S3 contain analogous histograms for the other cell line pairs. Each figure also includes the histogram corresponding to the box whose relocalization mean is the median over all identified boxes. Juxtaposed against the histograms are box relocalization means from the original PRIM analysis, thereby permitting readout of significant findings.

**Figure 3:**
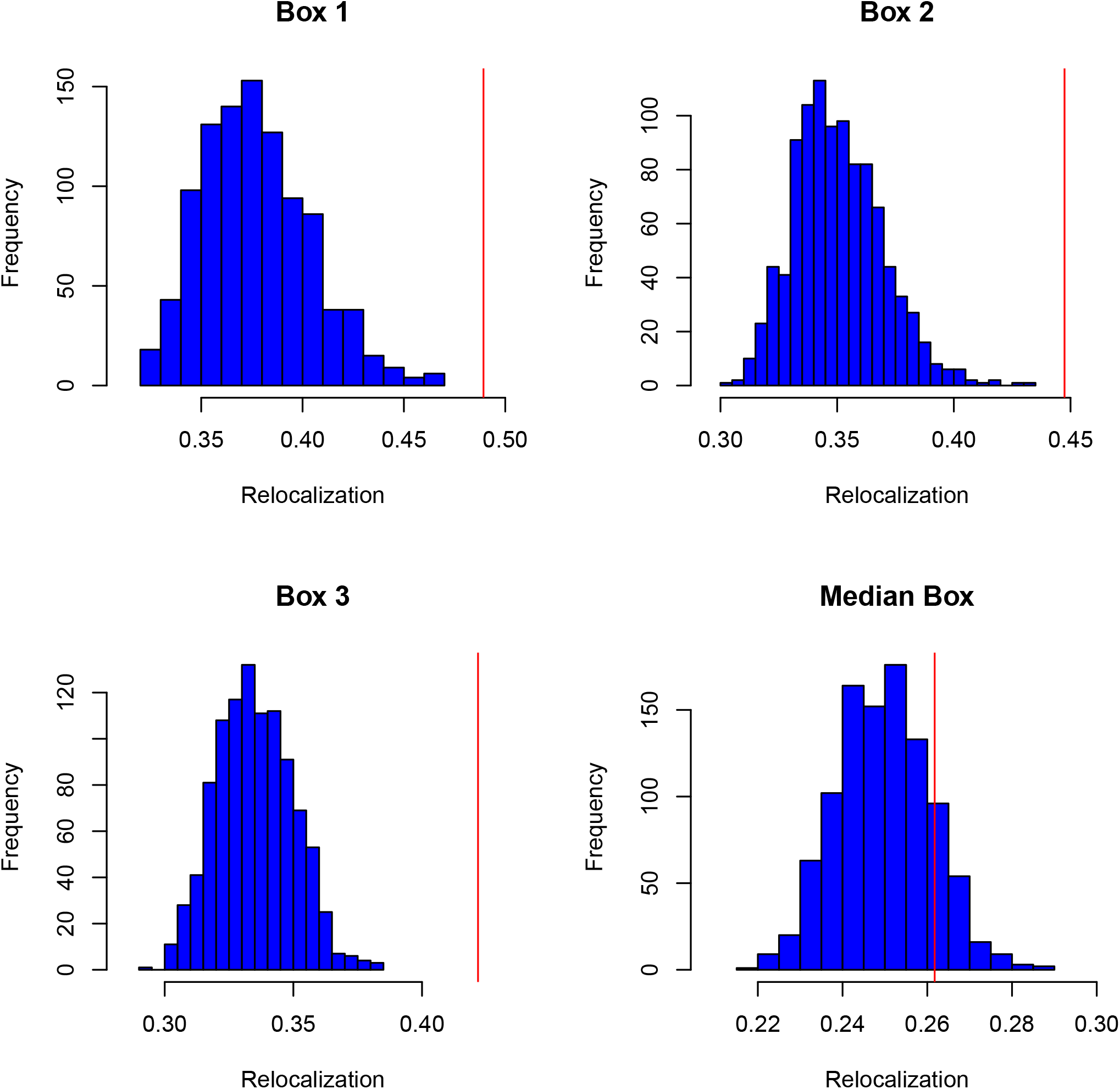
PRIM results for select (top 3, median) relocalization regions (boxes) between chromosome 21 of the multiMDS reconstructions of the **IMR90** cell line aligned to the **HMEC** cell line. The observed average relocalization distance for the respective boxes is indicated by the red vertical line while the blue histograms give null distributions obtained by permuting relocalizations over the aligned IMR90 reconstruction and re-applying the entire PRIM procedure 1000 times.

In both original and permuted PRIM applications we set *α*_*peel*_ = 10% and required that a box contain at least 0.1% of the total sample size. With these parameter settings the number of boxes extracted for the respective cell line comparisons were 31 (IMR90-HMEC), 34 (IMR90-HUVEC), 27 (HMEC-HUVEC) and 26 (K562-HUVEC). However, as suggested by the histograms corresponding to the median box, the number of significant boxes is appreciably less than these totals. While that number varies according to the cell line comparison, the region corresponding to the most extreme relocalizations (boxes 1 and 2) are significant throughout.

To place the significant regions detected in a genomic context, and to contrast with CWT-identified peaks, in Figure 4 we map the respective identified regions and segments to chromosome 21 genomic coordinates, overlaid on relocalization distance. Subsequently, in Figure 5, we position identified regions in their 3D context via superposition on the aligned 3D reconstruction.

**Figure 4:**
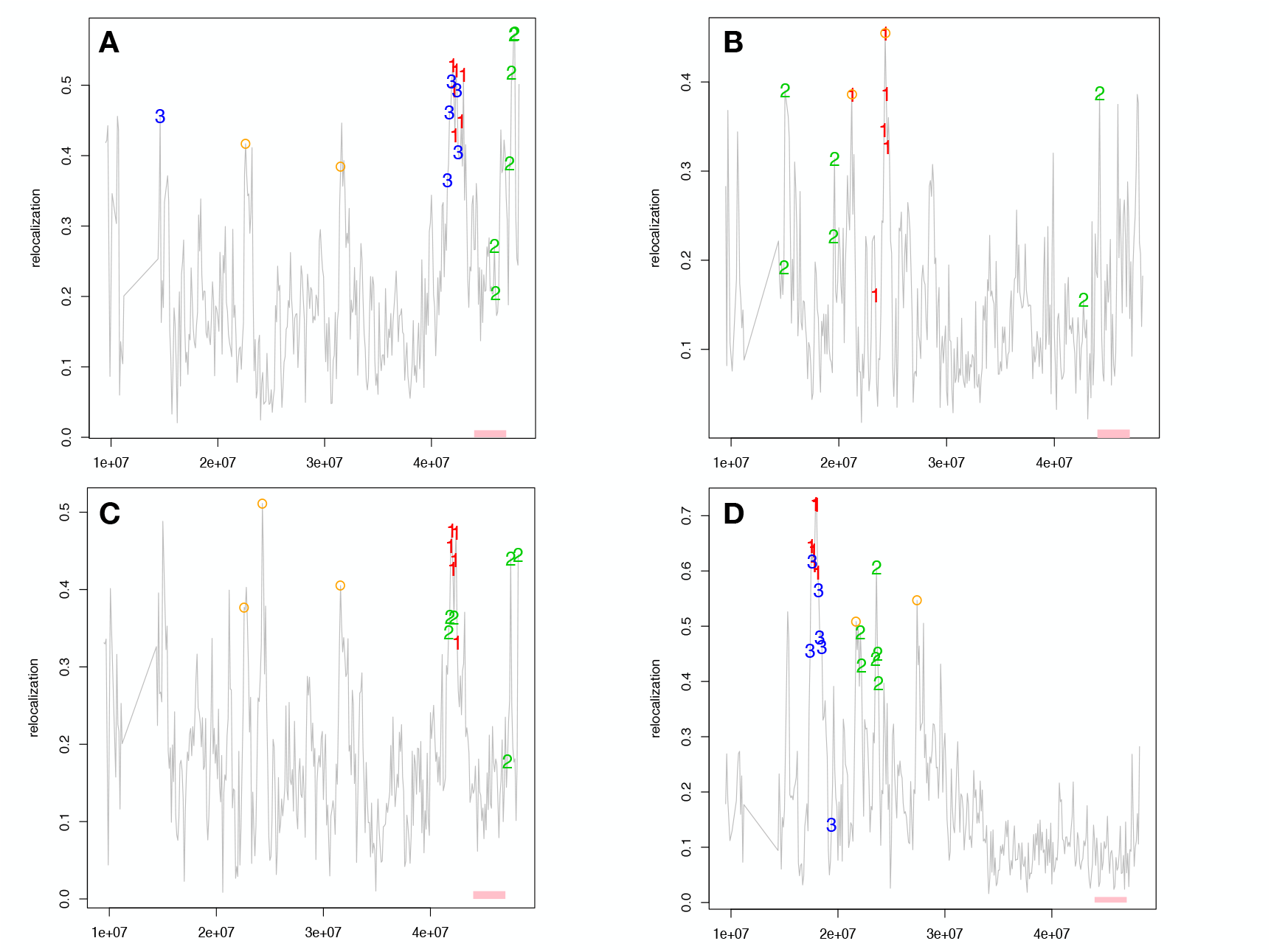
Relocalization distance between aligned multiMDS reconstructions as a function of chromosome 21 genomic cooordinates for A: IMR90 - HMEC, B: IMR90 - HUVEC, C: HMEC - HUVEC, and D: K562 - HUVEC cell line comparisons. In each panel loci belonging to significant PRIM-identified regions are shown (box 1: red, box 2: green, box 3: blue). The orange circles indicate loci of CWT-identified peaks and the pink bar corresponds to the key relocalization region (Rieber and Mahony (2019)).

**Figure 5:**
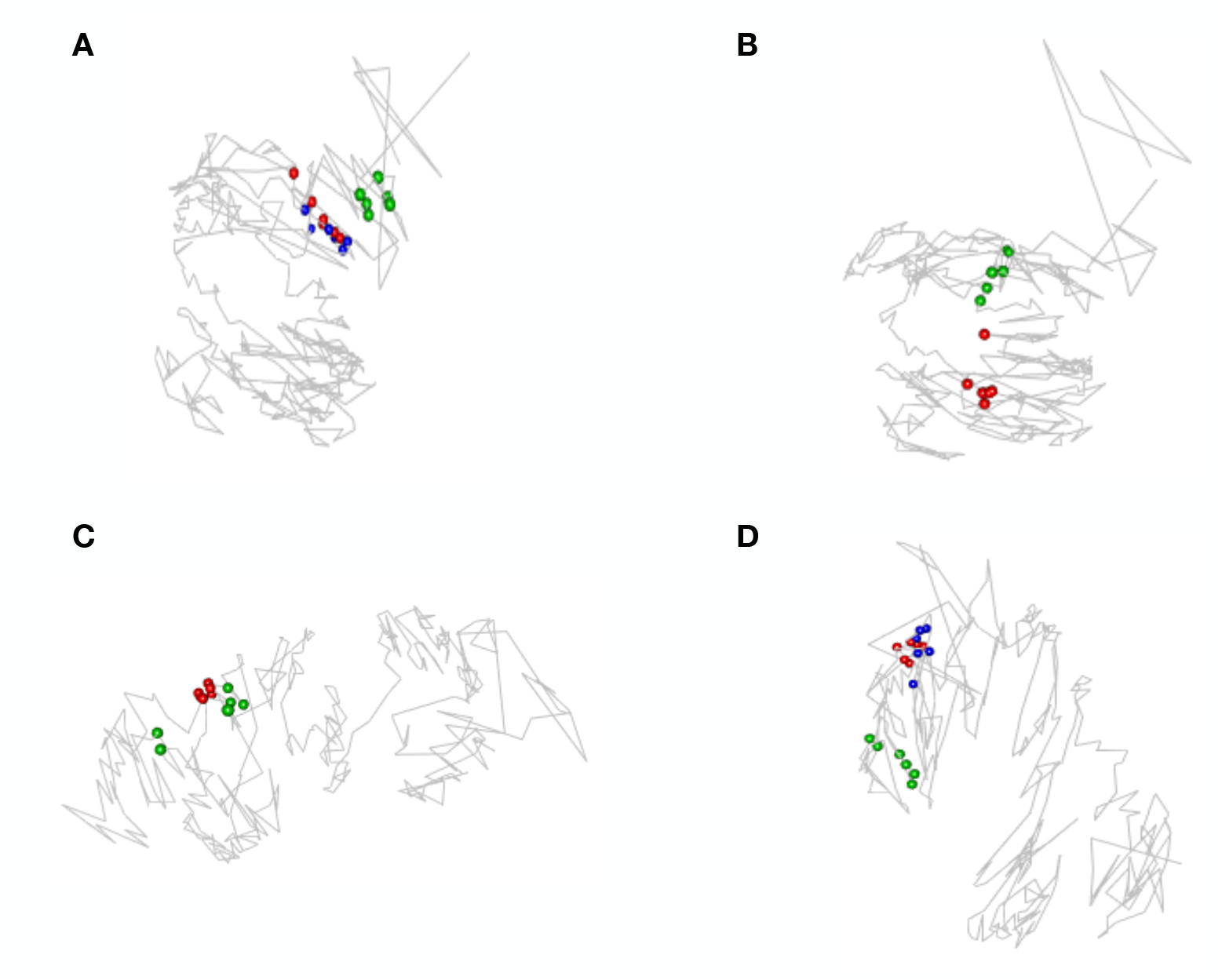
Positions of the significant PRIM boxes highlighted in the corresponding panels of Figure 4 superposed on the underlying multiMDS reconstructions: A: IMR90 - HMEC, B: IMR90 - HUVEC, C: HMEC - HUVEC, and D: K562 - HUVEC cell line comparisons.

Central to RM’s analysis is identification of a relocalization segment for IMR90, HMEC and HUVEC pairs at 47.4 - 47.5 Mb. However, identification of this key segment is not made via CWT peak finding, as evident from Figure 4 (orange circles). Rather, it follows visual inspection that is focused on segments for which differential (between cell types) compartment scores are insignificant which, as noted in Section 2.5, may be problematic. Importantly, relocalizations in this region are identified via PRIM (Figure 4), whereas the negative control comparison appropriately does not feature any such relocalization (Figure 4D). More generally, while the PRIM regions tend to be contiguous, there are instances of regions that are 3D but not 1D proximal, illustrating the value of pursuing a 3D relocalization identification scheme.

Results from interrogating significant regions and segments, in terms of epigenetic marker differences between the respective cell lines, are provided in in Section S4 of the supplementary material. The requirement that at least five marker values be available for each cell line meant that for several regions and segments limited the testing performed. Despite the modest sample sizes some consistent differences are evident with, for example, H3K4me1 having smaller signal for IMR90 than either HMEC of HUVEC at all significant regions or segments tested.

## 4 Discussion

This paper introduces PRIM as a technique for assessing chromatin relocalizations in 3D – reflecting the importance of 3D conformation in accord with the goals of 3D chromatin reconstruction – an improvement over existing 1D approaches. Relatedly, PRIM has been used to detect ‘3D hotspots’ of genomic features superposed on a 3D reconstruction (Capurso *and others* (2016)), analogous to superposition of relocalization distances. An important facet of such applications is that problem dimensions facilitate use of permutation-based inference in identifying significant regions. This follows from the covariate space dimension being limited to three (coordinate axes) and sample size being constrained by the resolution (number of bins) of the underlying Hi-C experiment which, with current technologies, are of order *∼*10^3^. For the analyses presented here, with 1000 permutations, CPU system times on a single processor 2.9 GHz machine are roughly 100 seconds. This contrasts with a PRIM investigation of identifying open clusters with extreme velocities in the Milky Way ((Segal and Segal, 2019)) where although, again, only three spatial dimensions were involved, the sample size (number of stars) was of order *∼*10^6^ resulting in constraints on permutation numbers, even after parallelizing over multiple processors.

However, PRIM analyses can also suffer from too few samples. RM ((Rieber and Mahony, 2019)) provide compelling applications of multiMDS in exploring relocalization differences between yeast species under varying galactose - glucose conditions. At the 32kb resolution used, bin numbers across the 16 chromosomes (with chromosome 12 partitioned at the nucleolus) range from 15 to 42. This is too sparse for effective use of PRIM: the designed emphasis on *patient* region elicitation requiring modest sample sizes. Looking ahead, the advent of much higher resolution conformation capture assays, for example CAP-C (You *and others* (2020)), may diminish this concern.

We have described refinements to multiMDS in Section 2.2. Similarly, the PRIM algorithm could be improved in several ways. Currently, the measure of regional (box) relocalization is simply the mean relocalization over constituent loci. Summaries that are both more robust and incorporate spread could be persued. While the R package utilized ((Duong, 2020)) features a tuning parameter that controls the minimum *mass* a candidate box must possess, a companion parameter controlling *volume* would preclude regions from becoming too spatially extensive. Devising approaches and/or guidelines for more systematic parameter tuning would be desirable.

Moving beyond PRIM-based methods for the task of eliciting 3D relocalization ‘hotspots’ deserves attention. Although use of PRIM with principle axes coordinate systems, with here the first component approximating the compartmental axis, have been advocated (Section 2.5), devising detection strategies that are invariant to 3D reconstruction orientation is a forefront need since, as noted in Section 2.5, results are not invariant to rotations of the 3D reconstruction yet reconstructions are inherently coordinate system-free. Here we have addressed this issue by (i) applying PRIM using a principal axes coordinate system and (ii) performing sensitivity analyses using numerous random structure rotations, but devising rotation invariant detection methods remains desirable. Developing such methods using topological data analysis techniques, including persistence homology ((Chazal *and others*, 2013)), is the subject of future work.

## 5 Software

An R function for performing PRIM-based detection of relocalization regions using PCA rotated 3D reconstructions and appraising significance via permutation is available from https://github.com/marksegal/prim-pca-permutation.

## Acknowledgments

The author thanks Lila Rieber and Shaun Mahony for graciously providing aligned multiMDS 3D reconstructions, Saurabh Asthana for obtaining ENCODE data, Elena Tuzhilina for performing reconstuction accuracy assessment and two referees for numerous helpful suggestions. *Conflict of Interest*: None declared.

