## Supplement for "Assessing chromatin relocalization in 3D using the patient rule induction method"

Mark R. Segal*

Department of Epidemiology and Biostatistics,
University of California, San Francisco CA 94143 USA

### S1 Concordance between compartmental and first principal component scores

| Cell Line 1 | Cell Line 2 | Number of Scores (Genomic Bins) | First PC Score : Compartment Score Correlation |
|---|---|---|---|
| IMR90 | HMEC | 347 | 0.88 |
| HMEC | IMR90 | 347 | 0.84 |
| IMR90 | HUVEC | 347 | 0.74 |
| HUVEC | IMR90 | 347 | 0.83 |
| HMEC | HUVEC | 348 | 0.77 |
| HUVEC | HMEC | 348 | 0.83 |
| K562 | HUVEC | 347 | 0.77 |
| HUVEC | K562 | 347 | 0.82 |

Table S1: Absolute correlations of first PC scores of multiMDS reconstruction of cell line 1 aligned to cell line 2 with the compartment score of cell line 1 as obtained from its Hi-C contact matrix.

### S2 Inter-cell Relocalization Region (box) Significance

Supplementary Figures S1 - S3 are analogous to Figure 3 of the main text. They show histograms of the permutation distribution – obtained by permuting (1000 times) the original relocalization distance values over the aligned structure coordinates – of the top three and median PRIM-selected relocalization distance region (box) means for IMR90 - HUVEC S1, HMEC - HUVEC S2, and K562 - HUVEC S3 cell line comparisons.

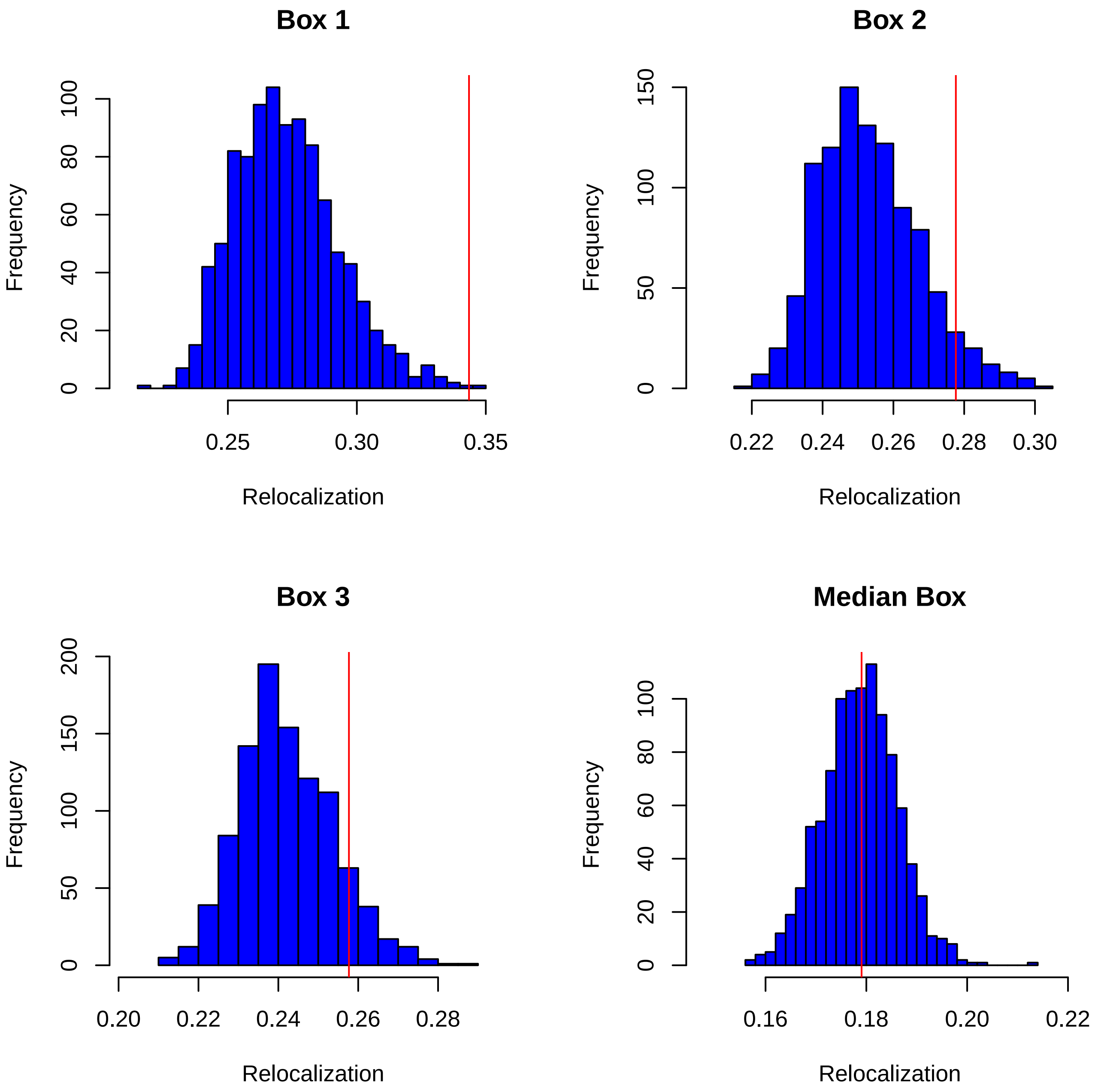


Figure S1: PRIM results for select (top 3, median) relocalization regions (boxes) between chromosome 21 of the multiMDS reconstructions of the **IMR90** cell line aligned to the **HUVEC** cell line. The observed average relocalization distance for the respective boxes is indicated by the red vertical line while the blue histograms give null distributions obtained by permuting relocalizations over the aligned IMR90 reconstruction and re-applying the entire PRIM procedure 1000 times.

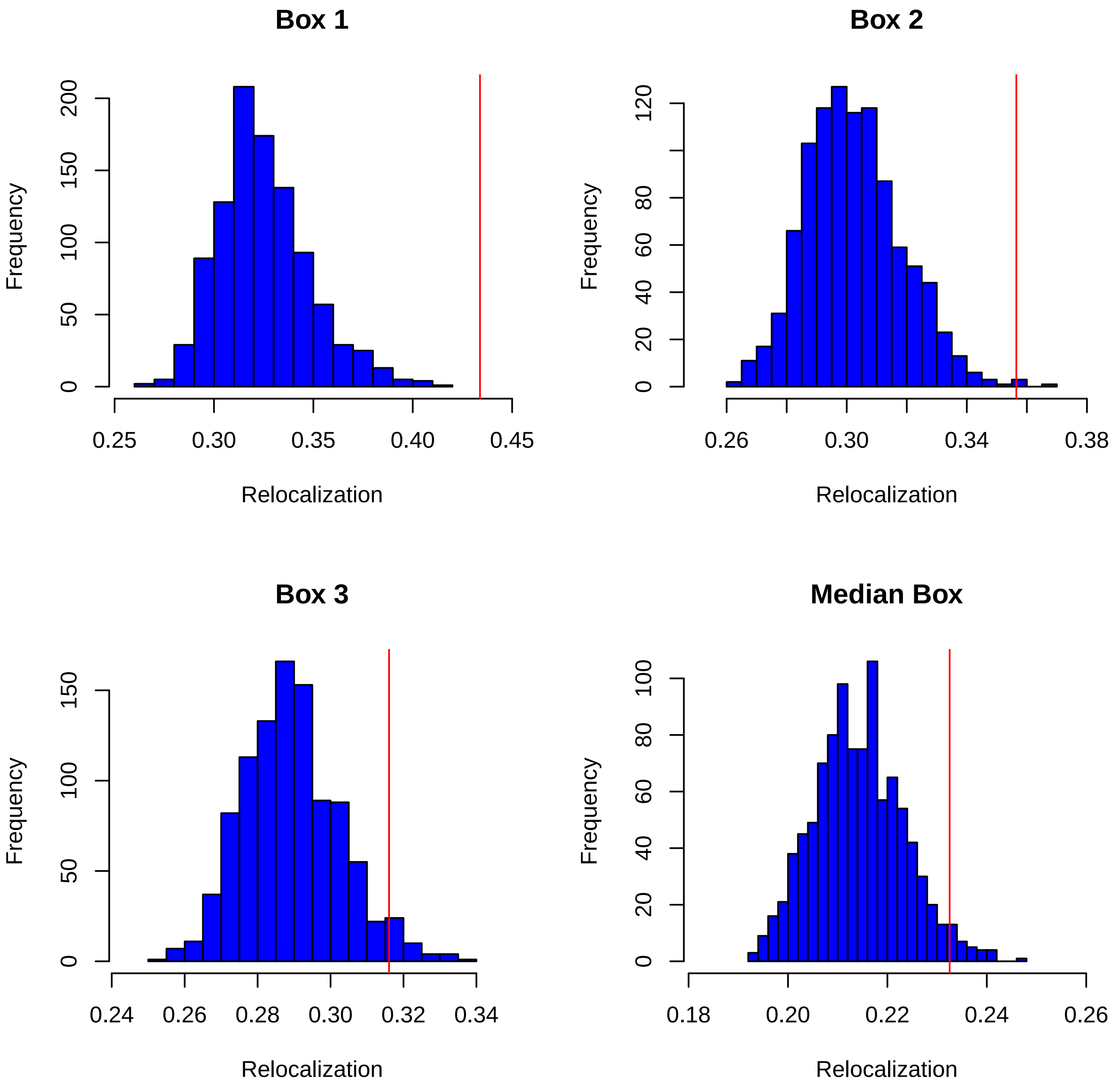


Figure S2: As for Figure S1 but showing PRIM relocalization regions (boxes) between chromosome 21 of the multiMDS reconstructions of the **HMEC** cell line aligned to the **HUVEC** cell line.

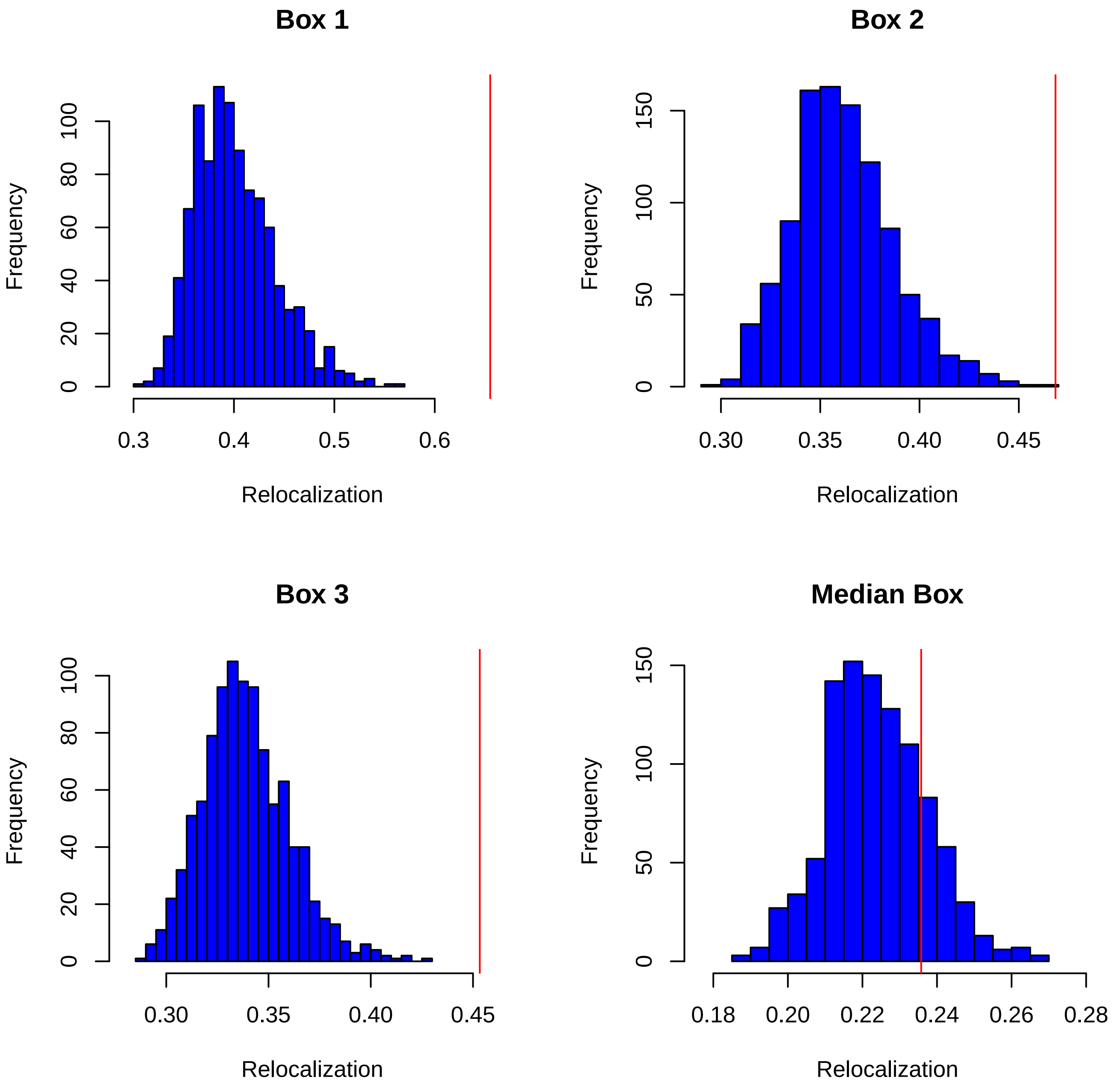


Figure S3: As for Figure S1 but showing PRIM relocalization regions (boxes) between chromosome 21 of the multiMDS reconstructions of the **K562** cell line aligned to the **HUVEC** cell line.

### S3 PRIM Region Identification Following Random Reconstruction Rotations

Supplementary Figures S4 - S7 are analogous to Figures 3 and 4 of the main text. Each figure displays results from two (out of 20) representative random rotations of an aligned, between cell line reconstruction pair. Note that since rotations are rigid, relocalization distance is preserved; i.e., Euclidean distances between original and rotated aligned reconstruction pairs are identical. Subsequent to applying the random rotation, instead of the rotation to principal axes that constituted our primary analyses, the same PRIM-based procedure for identifying relocalization regions is implemented. The only distinction from the approach presented in the main text (Sections 2.4 and 3.4; Figure 3, Supplementary Figures 1-3) is that the histograms depicting null reference distributions are based on 100 permutations instead of 1000, the reduction imposed solely to lessen the compute burden, as 20 differing random rotations, for each of the four between cell line comparisons, were conducted.

Synthesizing results over the complete set of random rotations leads to the following conclusions. There is generally good concordance among the regions identified by PRIM as significant relocalized over the random and principal axes rotated reconstructions. This agreement is best for IMR90-HMEC and K562-HUVEC comparisons, both in terms of region overlap and attendant significance. Nonetheless, there is notable variability in the *order* in which corresponding regions are selected.

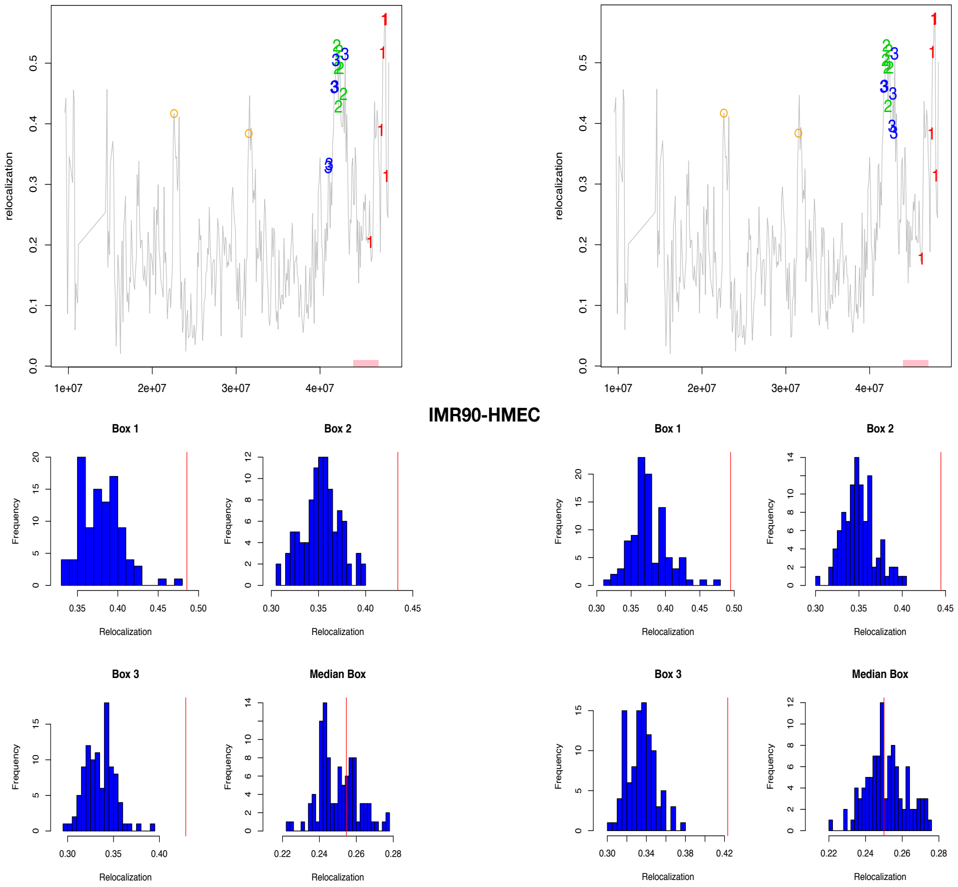


Figure S4: Top row: Relocalization distance between aligned multiMDS reconstructions as a function of chromosome 21 genomic cooordinates for the IMR90 - HMEC cell line comparisons. Each panel displays loci belonging to PRIM-identified regions (box 1: red, box 2: green, box 3: blue) following random rotation of the IMR90 reconstruction. The number of regions shown corresponds to the number of regions found significant in the original analysis (main text Figure 4). The orange circles indicate loci of CWT-identified peaks and the pink bar corresponds to the key relocalization region (Rieber and Mahony, 2019). Bottom row: Histograms (blue) showing null distributions obtained by permuting relocalization distances (from the panel above) over the rotated IMR90 reconstruction and re-applying the entire PRIM procedure 100 times, with results presented for the top 3 and median regions. The observed average relocalization distance for the boxes is indicated by the red vertical

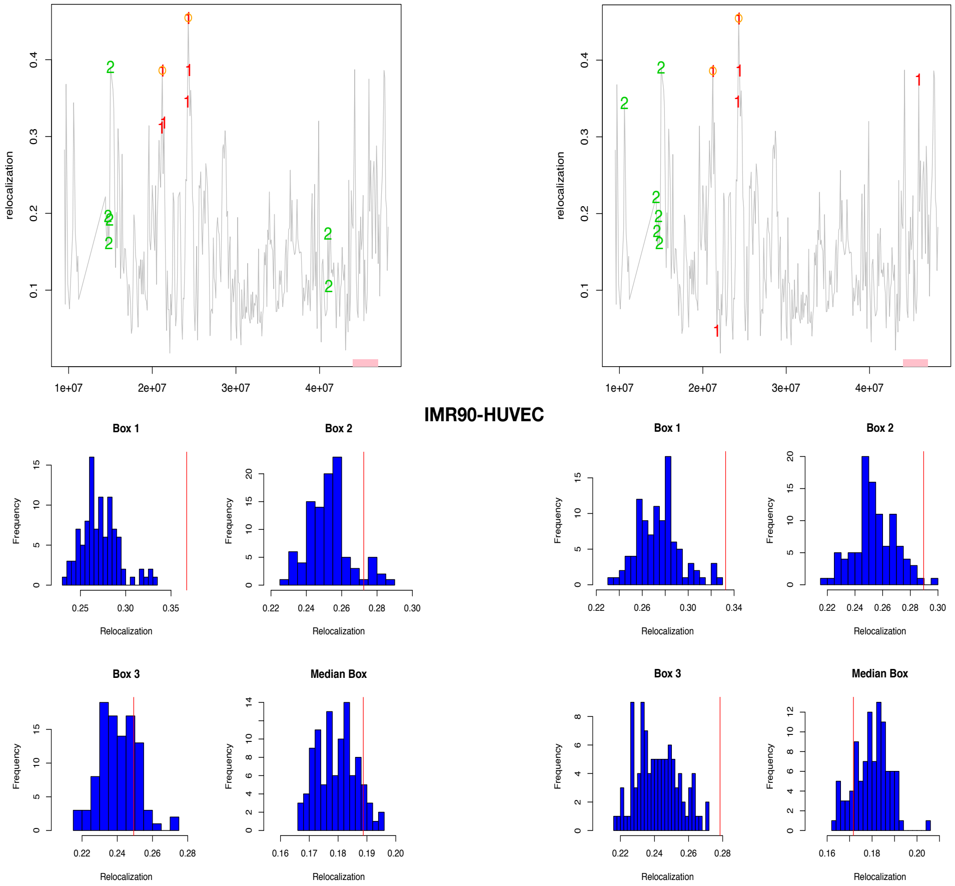


Figure S5: As for Figure S4 but showing PRIM relocalization regions (boxes) between chromosome 21 of the multiMDS reconstructions of the **HMEC** cell line aligned to the **HUVEC** cell line.

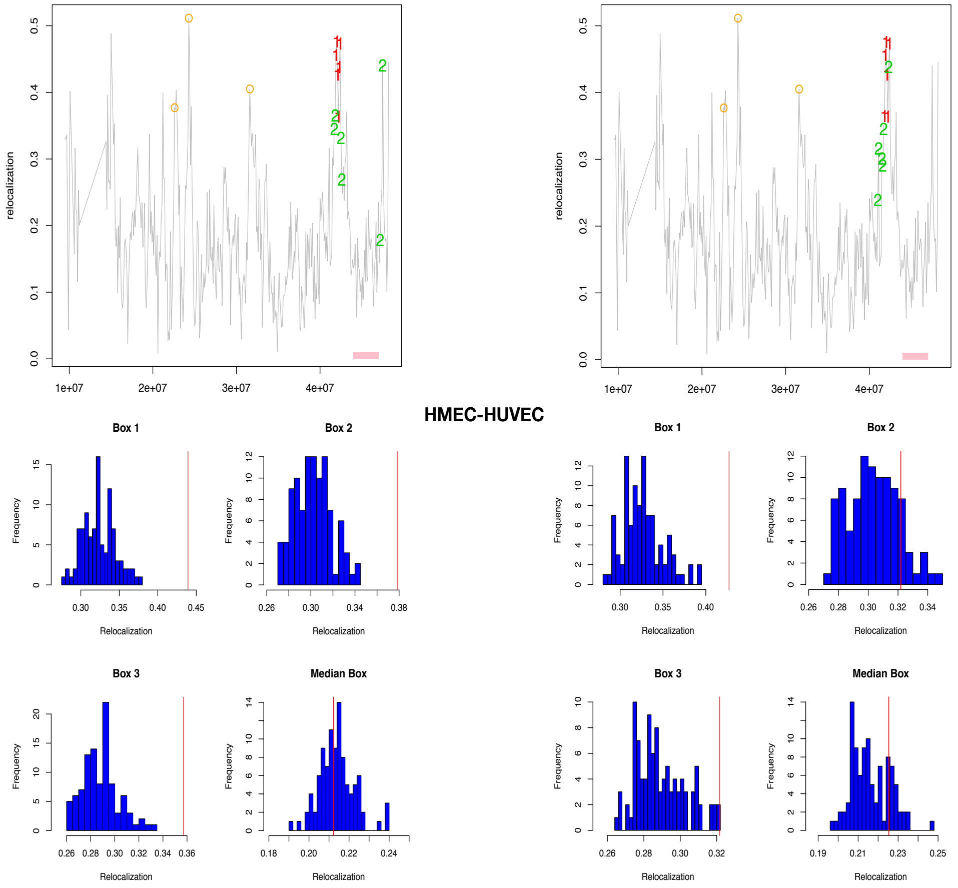


Figure S6: As for Figure S4 but showing PRIM relocalization regions (boxes) between chromosome 21 of the multiMDS reconstructions of the **K562** cell line aligned to the **HUVEC** cell line.

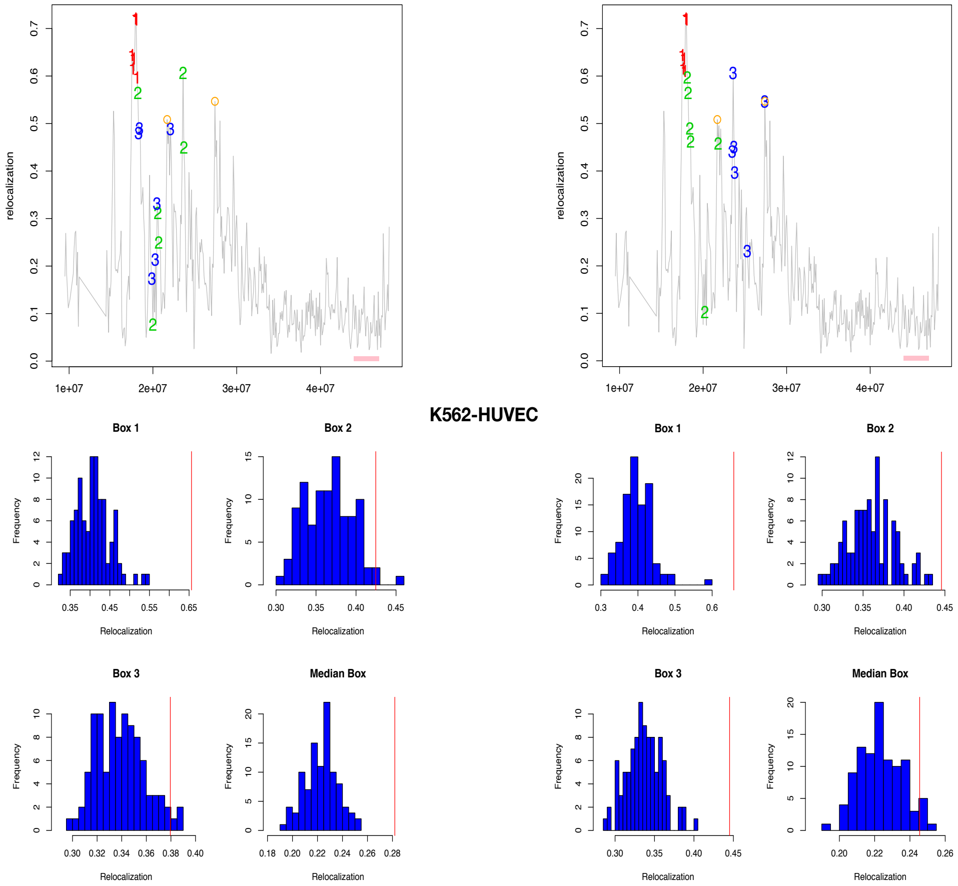


Figure S7: As for Figure S4 but showing PRIM relocalization regions (boxes) between chromosome 21 of the multiMDS reconstructions of the **K562** cell line aligned to the **HUVEC** cell line.

### S4 Chromatin marker differences in relocalization regions and segments

Having identified significant between cell line relocalization regions and segments (by PRIM and CWT respectively) we interrogated these (as recommended by two referees) for epigenetic modifications with a view to gaining putative insights into the drivers of the differing cell type configurations. However, while we examined a number of chromatin markers, this effort was limited by data sparsity. The epigenetic markers evaluated were H2A.Z, H3K27ac, H3K27me3, H3K36me3, H3K4me1, H3K4me2, H3K4me3 and H3K9ac and DNase accessibility, these being obtained from ENCODE as indicated in Section S4.2.

#### S4.1 Between cell line marker comparisons

Comparison between marker values in the significant relocalized regions / segments of the respective cell lines was effected by two sample t-tests, not prescribing equal variances and Welch modification to the degrees of freedom. Testing was subject to the requirement that each sample contain at least 5 marker values. This restriction precluded many comparisons, especially those involving HUVEC and K562 cell lines, which are omitted from the tabulated results that follow. A notable and consistent finding is that all comparisons involving significantly relocalized IMR90 regions feature significantly lower values of H3K4me1.

| Marker | t statistic | df | p-value |
|---|---|---|---|
| DNase | -1.613 | 20.08 | 1.22e-01 |
| H2A.Z | -4.125 | 54.18 | 1.28e-04 |
| H3K27ac | -2.031 | 28.59 | 5.16e-02 |
| H3K27me3 | 2.078 | 14.73 | 5.55e-02 |
| H3K36me3 | 2.023 | 44.26 | 4.90e-02 |
| H3K4me1 | -7.182 | 60.68 | 1.15e-09 |
| H3K4me2 | -3.608 | 40.35 | 8.39e-04 |
| H3K4me3 | -1.629 | 14.72 | 1.24e-01 |
| H3K9ac | -1.191 | 20.32 | 2.47e-01 |

Table S2: **IMR90 - HMEC**: significant PRIM region 1 (Figure 4A, red)

| Marker | t statistic | df | p-value |
|---|---|---|---|
| H2A.Z | -2.457 | 11.99 | 0.030 |
| H3K27me3 | -1.046 | 9.36 | 0.321 |
| H3K4me1 | -3.896 | 10.51 | 0.002 |
| H3K4me2 | -2.969 | 9.86 | 0.014 |

Table S3: **IMR90 - HMEC**: significant PRIM region 2 (Figure 4A, green)

| Marker | t statistic | df | p-value |
|---|---|---|---|
| DNase | -1.237 | 27.29 | 2.26e-01 |
| H2A.Z | -3.160 | 57.50 | 2.51e-03 |
| H3K27ac | -2.818 | 31.98 | 8.20e-03 |
| H3K27me3 | -0.021 | 14.23 | 9.82e-01 |
| H3K36me3 | 0.050 | 48.73 | 9.59e-01 |
| H3K4me1 | -4.980 | 62.50 | 5.29e-06 |
| H3K4me2 | -2.539 | 51.75 | 1.41e-02 |
| H3K4me3 | -1.168 | 8.97 | 2.72e-01 |

Table S4: **IMR90 - HMEC**: significant PRIM region 3 (Figure 4A, blue)

| Marker | t statistic | df | p-value |
|---|---|---|---|
| DNase | -4.675 | 15.06 | 2.95e-04 |
| H2A.Z | -3.876 | 38.51 | 4.01e-04 |
| H3K27ac | -2.569 | 13.69 | 2.25e-02 |
| H3K4me1 | -2.884 | 42.95 | 6.11e-03 |
| H3K4me2 | -1.835 | 25.31 | 7.81e-02 |
| H3K9ac | -1.336 | 14.77 | 2.01e-01 |

Table S5: **IMR90 - HMEC**: significant CWT segments (Figure 4A, orange circles ± 250kb)

| Marker | t statistic | df | p-value |
|---|---|---|---|
| DNase | 0.029 | 18.22 | 0.976 |

Table S6: **IMR90 - HUVEC**: significant PRIM region 1 (Figure 4B, red)

| Marker | t statistic | df | p-value |
|---|---|---|---|
| DNase | 1.242 | 51.38 | 0.219 |
| H2A.Z | -1.406 | 7.24 | 0.201 |
| H3K4me1 | -2.264 | 20.26 | 0.034 |

Table S7: **IMR90 - HUVEC**: significant PRIM region 2 (Figure 4B, green)

| Marker | t statistic | df | p-value |
|---|---|---|---|
| DNase | 0.955 | 57.40 | 0.343 |
| H2A.Z | 1.302 | 17.00 | 0.210 |
| H3K27ac | -1.067 | 14.69 | 0.302 |
| H3K27me3 | -1.313 | 30.82 | 0.198 |
| H3K36me3 | 1.833 | 12.57 | 0.091 |
| H3K4me1 | -3.259 | 42.40 | 0.002 |
| H3K4me2 | -1.524 | 32.25 | 0.137 |

Table S8: **IMR90 - HUVEC**: significant CWT segments (Figure 4B, orange circles ± 250kb)

| Marker | t statistic | df | p-value |
|---|---|---|---|
| DNase | 2.091 | 16.56 | 0.976 |
| H2A.Z | 2.660 | 57.95 | 0.010 |
| H3K27ac | 0.123 | 42.12 | 0.902 |
| H3K27me3 | -3.597 | 18.42 | 0.002 |
| H3K36me3 | -0.807 | 25.89 | 0.426 |
| H3K4me1 | -0.079 | 71.78 | 0.937 |
| H3K4me2 | 1.801 | 60.65 | 0.076 |
| H3K4me3 | 2.598 | 10.41 | 0.025 |
| H3K9ac | -1.877 | 29.00 | 0.070 |

Table S9: **HMEC - HUVEC**: significant PRIM region 1 (Figure 4C, red)

| Marker | t statistic | df | p-value |
|---|---|---|---|
| DNase | 3.669 | 13.22 | 0.002 |
| H2A.Z | 3.338 | 44.94 | 0.001 |
| H3K27ac | 0.574 | 17.93 | 0.572 |
| H3K4me1 | 0.941 | 20.24 | 0.357 |
| H3K4me2 | 0.330 | 10.56 | 0.747 |

Table S10: **HMEC - HUVEC**: significant PRIM region 2 (Figure 4C, green)

| Marker | t statistic | df | p-value |
|---|---|---|---|
| DNase | 4.462 | 11.37 | 0.001 |
| H3K4me1 | -0.096 | 16.06 | 0.924 |
| H3K4me2 | -0.018 | 19.86 | 0.985 |

Table S11: **HMEC - HUVEC**: significant CWT segments (Figure 4C, orange circles ± 250kb)

| Marker | t statistic | df | p-value |
|---|---|---|---|
| DNase | 0.607 | 35.28 | 0.547 |
| H2A.Z | 1.304 | 22.10 | 0.205 |
| H3K27me3 | -0.568. | 9.89 | 0.582 |
| H3K4me2 | 0.232 | 19.95 | 0.818 |

Table S12: **K562 - HUVEC**: significant PRIM region 1 (Figure 4D, red)

| Marker | t statistic | df | p-value |
|---|---|---|---|
| DNase | 1.457 | 15.40 | 0.165 |

Table S13: **K562 - HUVEC**: significant PRIM region 3 (Figure 4D, BLUE)

| Marker | t statistic | df | p-value |
|---|---|---|---|
| DNase | 1.060 | 39.32 | 0.295 |

Table S14: **K562 - HUVEC**: significant CWT segments (Figure 4D, orange circles ± 250kb)

#### S4.2 Sources for ENCODE chromatin modification marks

| Cell Line | Marker | Source url: `https://www.encodeproject.org/files/...` |
|---|---|---|
| IMR90 | H2A.Z | `ENCFF961ORU/@@download/ENCFF961ORU.bed.gz` |
| IMR90 | H3K4me1 | `ENCFF611UWF/@@download/ENCFF611UWF.bed.gz` |
| IMR90 | H3K4me2 | `ENCFF438DUR/@@download/ENCFF438DUR.bed.gz` |
| IMR90 | H3K4me3 | `ENCFF093NQC/@@download/ENCFF093NQC.bed.gz` |
| IMR90 | H3K9ac | `ENCFF309ZMM/@@download/ENCFF309ZMM.bed.gz` |
| IMR90 | H3K27ac | `ENCFF805GNH/@@download/ENCFF805GNH.bed.gz` |
| IMR90 | H3K27me3 | `ENCFF336IXL/@@download/ENCFF336IXL.bed.gz` |
| IMR90 | H3K36me3 | `ENCFF449ADN/@@download/ENCFF449ADN.bed.gz` |
| IMR90 | DNase | `ENCFF221SZF/@@download/ENCFF221SZF.bed.gz` |
| HMEC | H2A.Z | `ENCFF360HTG/@@download/ENCFF360HTG.bed.gz` |
| HMEC | H3K4me1 | `ENCFF401CLZ/@@download/ENCFF401CLZ.bed.gz` |
| HMEC | H3K4me2 | `ENCFF787CWV/@@download/ENCFF787CWV.bed.gz` |
| HMEC | H3K4me3 | `ENCFF898NGW/@@download/ENCFF898NGW.bed.gz` |
| HMEC | H3K9ac | `ENCFF152HYX/@@download/ENCFF152HYX.bed.gz` |
| HMEC | H3K27ac | `ENCFF929WUQ/@@download/ENCFF929WUQ.bed.gz` |
| HMEC | H3K27me3 | `ENCFF518UWB/@@download/ENCFF518UWB.bed.gz` |
| HMEC | H3K36me3 | `ENCFF745BUS/@@download/ENCFF745BUS.bed.gz` |
| HMEC | DNase | `ENCFF628QME/@@download/ENCFF628QME.bed.gz` |
| HUVEC | H2A.Z | `ENCFF088TGM/@@download/ENCFF088TGM.bed.gz` |
| HUVEC | H3K4me1 | `ENCFF213BAF/@@download/ENCFF213BAF.bed.gz` |
| HUVEC | H3K4me2 | `ENCFF882WZG/@@download/ENCFF882WZG.bed.gz` |
| HUVEC | H3K4me3 | `ENCFF524QAC/@@download/ENCFF524QAC.bed.gz` |
| HUVEC | H3K9ac | `ENCFF568IPQ/@@download/ENCFF568IPQ.bed.gz` |
| HUVEC | H3K27ac | `ENCFF077LGZ/@@download/ENCFF077LGZ.bed.gz` |
| HUVEC | H3K27me3 | `ENCFF977TUS/@@download/ENCFF977TUS.bed.gz` |
| HUVEC | H3K36me3 | `ENCFF200RLK/@@download/ENCFF200RLK.bed.gz` |
| HUVEC | DNase | `ENCFF080VVT/@@download/ENCFF080VVT.bed.gz` |
| K562 | H2A.Z | `ENCFF213OTI/@@download/ENCFF213OTI.bed.gz` |
| K562 | H3K4me1 | `ENCFF759NWD/@@download/ENCFF759NWD.bed.gz` |
| K562 | H3K4me2 | `ENCFF749KLQ/@@download/ENCFF749KLQ.bed.gz` |
| K562 | H3K4me3 | `ENCFF706WUF/@@download/ENCFF706WUF.bed.gz` |
| K562 | H3K9ac | `ENCFF891CHI/@@download/ENCFF891CHI.bed.gz` |
| K562 | H3K27ac | `ENCFF864OSZ/@@download/ENCFF864OSZ.bed.gz` |
| K562 | H3K27me3 | `ENCFF801AHF/@@download/ENCFF801AHF.bed.gz` |
| K562 | H3K36me3 | `ENCFF561OUZ/@@download/ENCFF561OUZ.bed.gz` |
| K562 | DNase | `ENCFF185XRG/@@download/ENCFF185XRG.bed.gz` |

Table S15: Data sources for ENCODE chromatin markers for the four comparison cell lines.
